# Practicing one thing at a time: the secret to reward-based learning?

**DOI:** 10.1101/745778

**Authors:** Katinka van der Kooij, Nina M van Mastrigt, Jeroen BJ Smeets

## Abstract

Binary reward feedback on movement success is sufficient for learning in some simple reaching tasks, but not in some more complex ones. It is unclear what the critical conditions for learning are. Here, we ask how reward-based sensorimotor learning depends on the number of factors that are task-relevant. In a task that involves two factors, we test whether learning improves by giving feedback on each factor in a separate phase of the learning. Participants learned to perform a 3D trajectory matching task on the basis of binary reward-feedback in three phases. In the first and second phase, the reward could be based on the produced slant, the produced length or the combination of the two. In the third phase, the feedback was always based on the combination of the two factors. The results showed that reward-based learning did not depend on the number of factors that were task-relevant. Consistently, providing feedback on a single factor in the first two phases did not improve motor learning in the third phase.

## Introduction

Katy practices a dance move while her trainer tells her whether her attempts are successful or not. Each time she hears that she was successful, a reward signal is delivered to her brain that can form the basis for motor learning ^1,2^. Several mechanisms have been proposed to underlie this learning. For instance, Katy may directly form associations between actions and success which would result in action-specific learning ^1,3^. Another possibility is that she may learn from successful exploration ^4–6^, which could potentially transfer across actions. The type of mechanism that we focus on in this study is a learning mechanism that relies on the exploitation of successful exploration.

The task in which reward-based learning has been typically tested is a far cry from the complexity of ‘real-life’ tasks such as Katy’s dance training ^7^. In the commonly used ‘visuomotor rotation’ paradigm ^1,4–6,8,9^ movements are in a horizontal plane in which the relation between visual and motor direction is rotated. In this paradigm, participants make center-out reaching movements in which reward feedback is based on a single factor: the angular error. In most natural tasks, in contrast, many factors are task-relevant. For instance, Katy’s dance move will only be successful if she gets both the position and timing right. We previously showed that learning of a visuomotor perturbation in a three-dimensional pointing task was not possible when participants received feedback based on a three-dimensional position error ^10–12^. However, learning did occur when the feedback was based on the perturbed dimension only ^10^. This suggests that reward-based learning depends on the number of factors that are task-relevant.

In the current study, we test two hypotheses. The first hypothesis is that learning of a factor improves when it is the only task-relevant factor. The second hypothesis is that learning of multiple factors improves by factorizing the feedback (giving feedback on each factor in a separate phase of the learning). We test these hypotheses in a three-dimensional trajectory matching task akin to trajectory learning tasks used in other studies on reward-based learning ^3,13^. We asked participants to copy a remembered simple trajectory – a slanted line – by moving the unseen hand. Without training, participants make systematic errors in this task, and the aim of the feedback was to reduce these errors. Feedback could be based on errors in produced slant, errors in produced length or the combination of these error components (combined feedback).

We divided the participants in three groups that differed in the way they received feedback during the three learning phases of the experiment. A ‘Slant First’ group received feedback on slant in a first learning phase, feedback on length in a second learning phase and ended with combined feedback. A ‘Length First’ group received feedback on length in their first learning phase, feedback on slant in their second learning phase and ended with combined feedback. A ‘Combined’ group, finally, received combined feedback throughout all learning phases of the experiment. If reward-based learning depends on the number of factors that are task-relevant, learning of a single factor would be faster when it is the only factor that is task-relevant. Furthermore, if factorized feedback improves reward-based learning, we predict that at the end of the third learning phase, the combined error is reduced more in the Length First and Slant First groups compared to the Combined group. In addition to testing these predictions, we assessed how exploration depends on the feedback.

## Results

In virtual reality, participants viewed a line slanted in the sagittal plane, and copied this line with an invisible handheld controller. They did so in five phases of 50 trials each. In a baseline and retention phase, no performance feedback was provided. In the three intermediate learning phases, binary score reward feedback was provided. As expected, participants showed biases in both slant and length that tended to be reduced in the third learning phase (Figure 1a). Because motivation may affect how participants learn from score reward, we asked participants about their motivation following each phase. Reported motivation in the three groups was similar (Figure 1b). Therefore, between-group differences in learning should probably not be attributed to differences in motivation.

**Figure 1.**
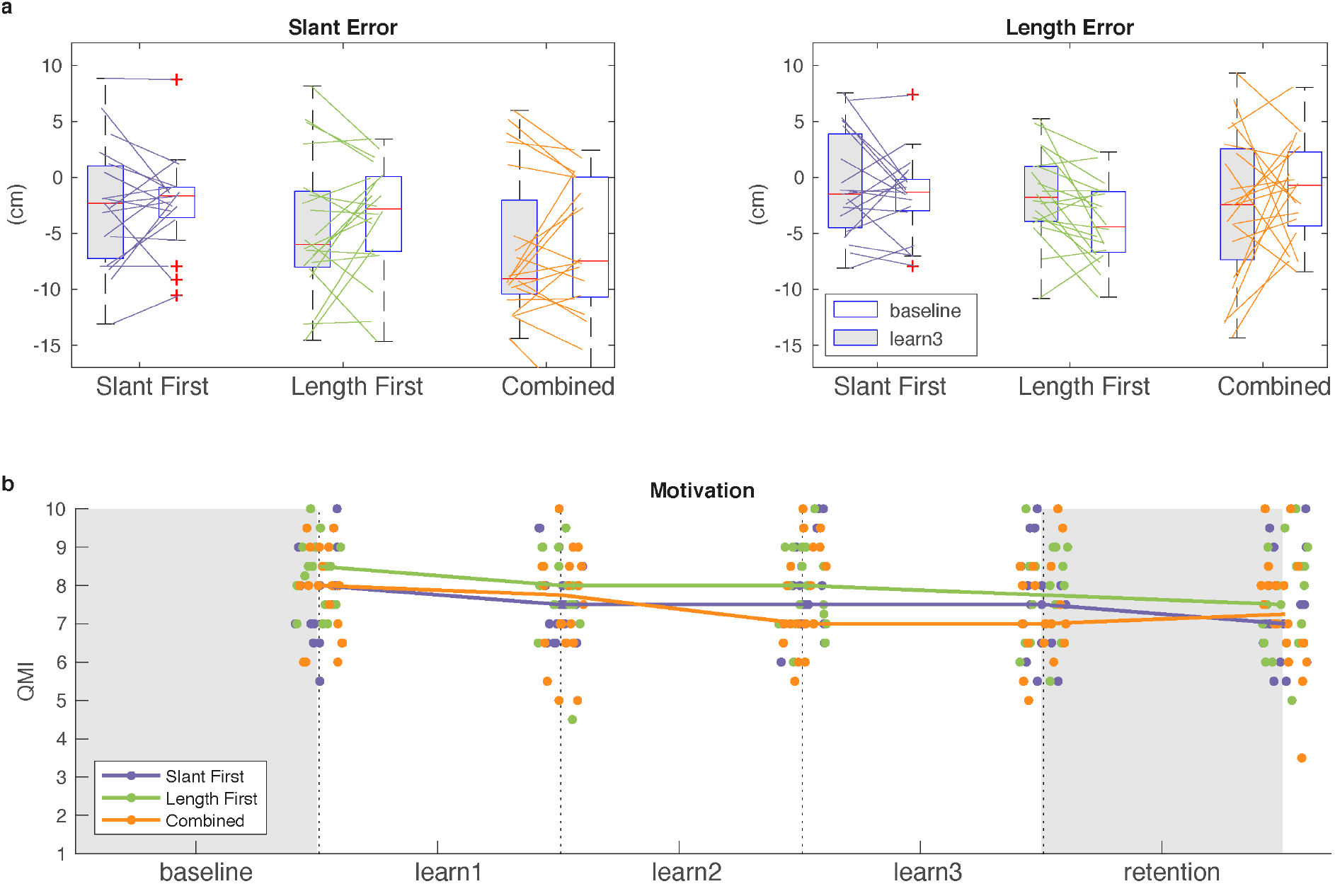
Overview of individual performance and motivation. **a)** Slant error (e_θ_; left panel) and length error (e_l_; right panel) in the baseline phase and in the final learning phase. **b**) Motivation of the individuals (dots) in each group with their median (lines) as a function of experimental phase. We applied a small horizontal scatter to the dots to show the individual data.

To study learning, we were interested in how the errors change relative to baseline. We therefore used the normalized error, which has a value of one at baseline and is zero if learning is complete. Overall, the normalized error in slant, length and the combined factor was reduced in the third learning phase and this learning was at least partially retained in the retention phase (Figure 2a).

**Figure 2.**
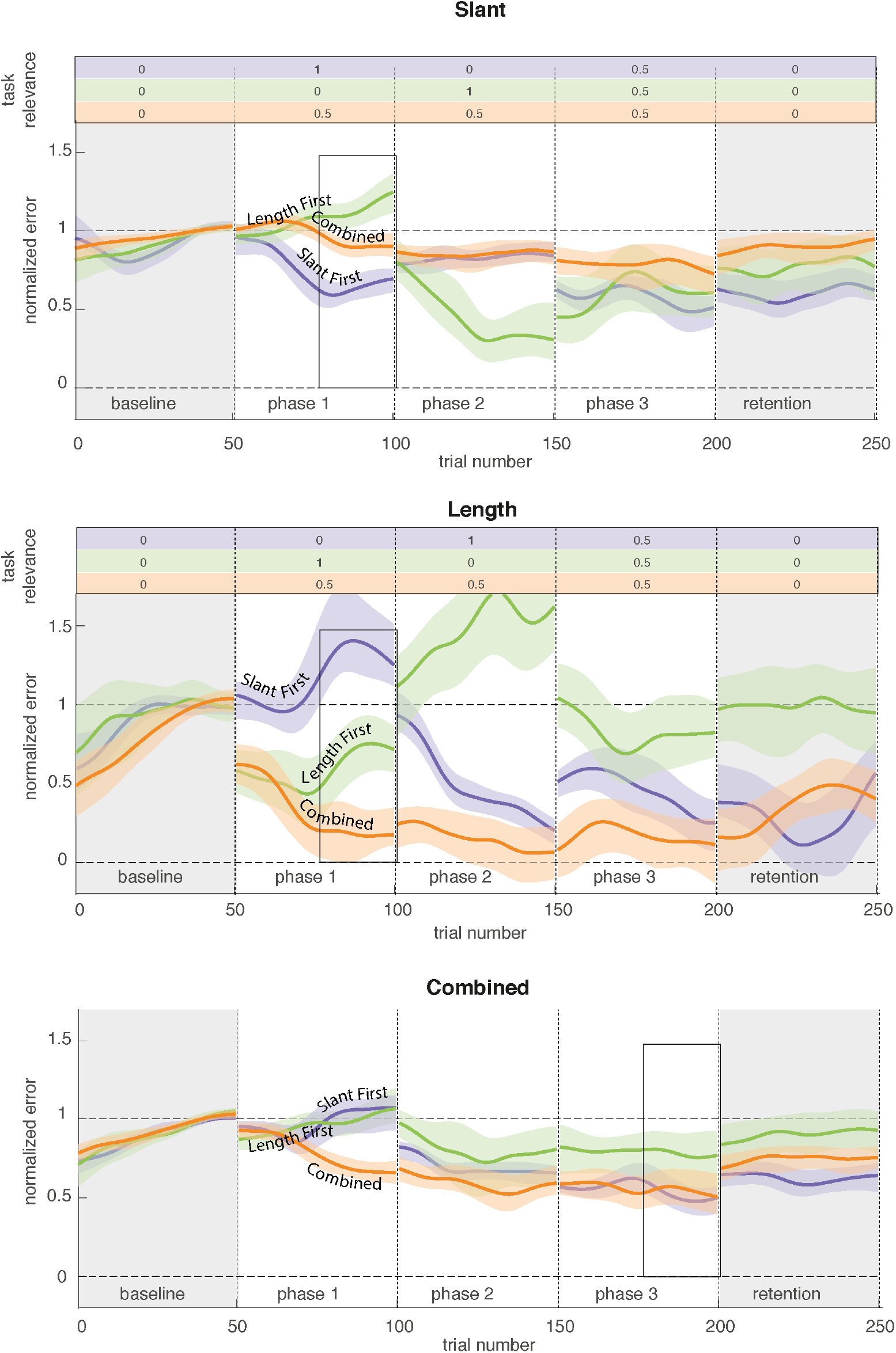
Results on learning. Median normalized error with 95% confidence interval as a function of trial number for the three groups. Data are smoothed with a Gaussian kernel (σ = 5 trials) for each phase separately. The dashed horizontal lines indicate the value that would indicate no learning (1) and complete learning (0). Grey background indicates phases without feedback. Rectangles indicate the episode within which the learning was analyzed.

We first tested whether learning of a single factor improves when it is more task-relevant. For task relevance, we take a value of 1 when a factor is the only task-relevant factor, a value of 0.5 when there is one other task-relevant factor and 0 was the factor is not task-relevant. As a measure of learning, we analyzed the fraction of the baseline error that was removed in the last 20 trials of each phase (the ‘gain’). In contrast to our prediction, learning the in the first learning phase was not better for factorized learning than for combined learning. This was the case both for slant (Slant First group with task-relevance = 1, compared to the Combined group with task-relevance = 0.5; Medians 0.51 and 0.37, *z* = 0.13, *p* = 0.45) as well as for length (Length First group with task-relevance = 1 compared to the Combined group with task-relevance = 0.5; Medians 0.34 and 0.85, *z* = −2.02, *p* = 0.98).

Next, we analyzed whether factorized feedback improved learning of the combined factor at the end of the third learning phase (Figure 2c). We focused on the third learning phase rather than on the retention phase because our primary interest is in learning, not retention. In contrast to the prediction that the groups that had received factorized feedback would have learned more than the Combined group, learning of the combined factor in the Slant First group (median = 0.54) or Length First group (median = 0.17) was not better (*z* = 0.41, *p* = 0.34 and *z* = −1.18, *p* = 0.88, respectively) than for the Combined group (median = 0.44). Thus we found that participants could learn the task and that learning was not improved by factorized feedback. In the next paragraphs we address exploration.

Because theories propose that reward-based learning relies on exploration ^3,4,6^, we assessed how exploration depends on the feedback and task relevance. Exploration will increase trial-by-trial changes. Therefore we analyze exploration from trial-by-trial changes. Trial-by-trial changes (*Δ*) were calculated as the amplitude of each change in the normalized error from trial *t*-1 to trial *t*. Several studies have found that trial-by-trial changes are larger following non-rewarded trials than following rewarded trials ^4,5,10,12,14,15^. To test whether this effect existed in the current study, we calculated the median trial-by-trial change in the combined error (*Δ_Combined_*) following non-rewarded trials and following rewarded trials within the three learning phases. A one-sided Wilcoxon sign rank test showed that, as expected, changes were larger following non-rewarded trials: medians 2.05 cm and 1.38 cm, *z* = 6.56, *p* < 0.01 (Figure 3a).

**Figure 3.**
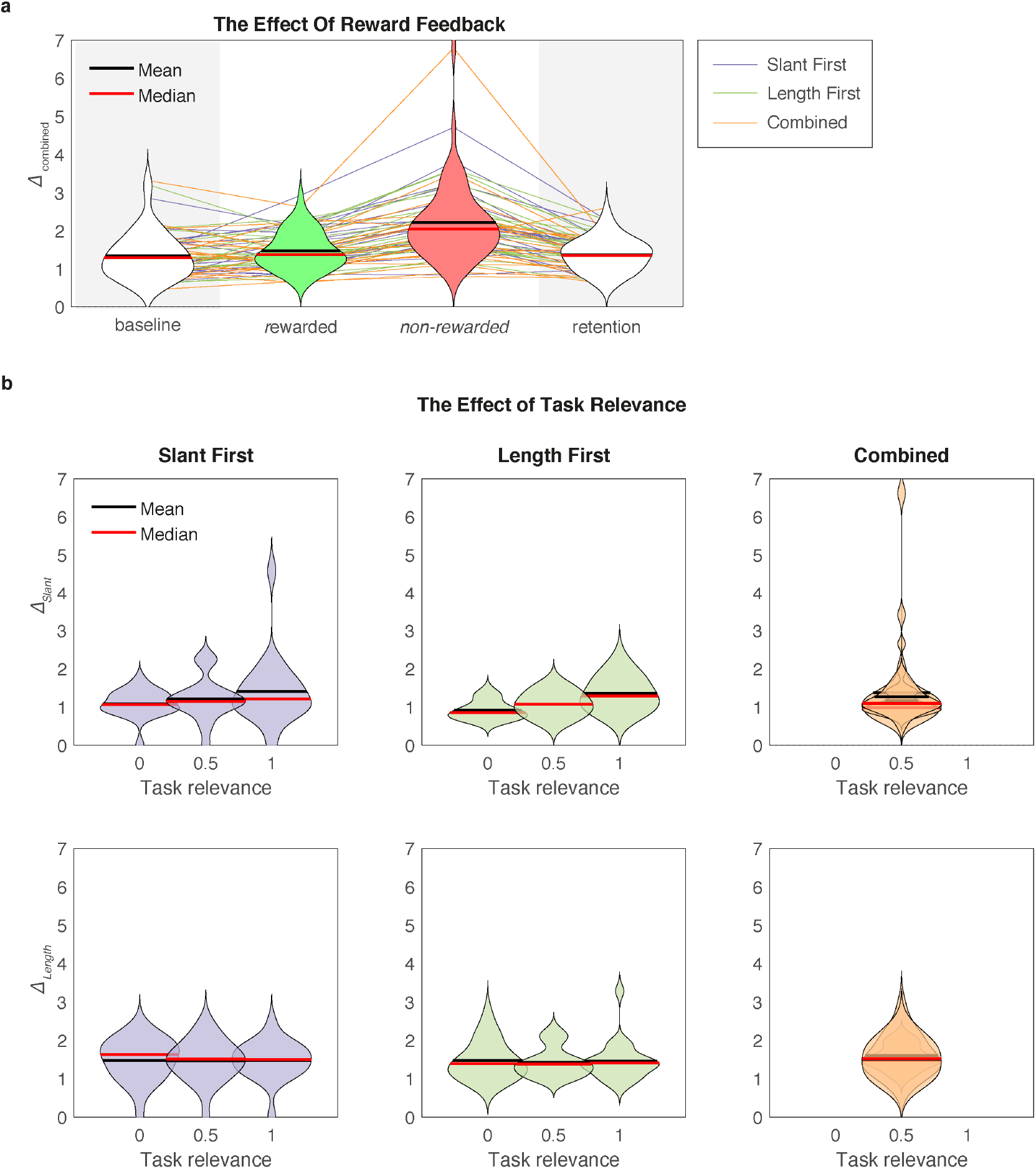
Violin plots ^16^ of the effect of reward and task-relevance on exploration. **a)** Trial-by-trial changes in the combined error (Δ_Combined_) following rewarded and non-rewarded trials in the learning phases and in the phases without feedback (grey background), **b)** Trial-by-trial changes in the slant error (Δ_Slant_) and in the length error (Δ_Length_) as a function of task relevance within the three groups. For the Combined group the task relevance was 0.5 in all three phases. Therefore, the data from the three phases overlap. Violins have been smoothed with default MatLab kernel density smoothing, depending on the number of points and range of the data.

Next, we tested whether a factor is explored more when it is more task-relevant. To this end, we performed a linear regression on the trial-by-trial variation in slant and length (*Δ_Slant_* and *Δ_Length_)* as a function of the factors’ task relevance and phase (1,2,3). Phase was included in this analysis because exploration might decrease with phase, possibly confounding the effect of task-relevance on trial-by-trial variation. As the value for task relevance we used 0 when no feedback was provided on the factor, 1 when it was the only factor that feedback was provided on and 0.5 when the feedback was provided on the combined error. For slant, we found that trial-by-trial variation depended positively on task relevance (*slope* = 0.32 cm; 95% CI: [0.04, 0.60]) but not on phase (*slope* = −0.02 cm; CI [−0.14, 0.09]). For length, in contrast, we did not find that trial-by-trial variation depended on the task-relevance (*slope* = 0.02 cm; CI [−0.18, 0.21]), neither did the trial-by-trial variation in length depend on phase (slope = −0.01 cm; CI [−0.09, 0.07]).

To test whether these results were specific to the case in which task-relevance was explicitly defined in the instructions, we analyzed the Combined group in which task relevance of slant and length was not explicitly defined but implicitly determined by the relative amplitude of slant and length biases. We therefore calculated the ‘implicit task-relevance’ for a factor as the bias amplitude divided by the sum of the bias amplitude in slant and length. Next, we performed a rank order correlation between the implicit task relevance for a factor and the trial-by-trial changes of this factor. There was no significant correlation (for slant: *R* = 0.03, *p* = 0.80; for length: *R* = 0.02, *p* = 0.86).

The explorative exit questionnaire on explicit strategies provided no additional insights.

## Discussion

In this study, we investigated whether reward-based motor learning depends on whether one or two factors are task-relevant. In addition, we tested whether factorizing feedback on two factors improved learning of these factors. We addressed these questions in a novel 3D trajectory matching task that allowed us to test motor learning without perturbing feedback to impose errors. For both slant and length, we found that unique task relevance of a factor did not improve learning of that factor. Consistently, factorizing feedback did not improve learning of the combination of the factors: participants who received feedback on slant and length at the same time learned the task equally well as participants who received factorized feedback. Below we discuss what these results imply for the mechanisms of reward-based learning.

In the present study, learning did not depend on the number of factors that are task-relevant. This seems in contrast with our earlier finding that learning of a lateral perturbation on reaching was possible with feedback on one factor (the lateral dimension), but not if feedback was based on two additional factors (three-dimensional position; three-dimensional position; ^10^. Three differences in the experimental design may underlie the difference in results. First, the number of factors to be learned was smaller in the current study: 2 instead of 3, so the limitation could be on learning three factors. The second difference is related to the noise in the task-relevant factors. Noise hampers learning ^5^ and an additional factor may hamper learning by adding noise to the signal that the reward is based on. In van der Kooij & Smeets (2019) we studied the addition of depth and elevation to a lateral task. Depth perception is associated with a higher level of perceptual noise than perception of lateral position-especially in a virtual reality setup ^17^. Therefore, adding a depth factor may have added a significant amount of noise to the motor output that the reward was based on. The factors in the current study both depend on depth perception and might have been associated with more comparable levels of perceptual noise. Hence, learning of the 3D task in our previous study may have been impaired by perceptual noise rather than by an inability to learn more than one factor in parallel. The third difference is related to what participants had to learn. In the previous study, a visual perturbation was used to impose errors whereas in the current study no perturbation was used. Although a visual perturbation defines movements that miss the target as successful, participants may proprioceptively sense that the movement missed the target ^18^. Such proprioceptive errors could have interfered with learning from the reward in our previous study, similar to how reward can interfere with learning from error ^12^. To conclude, we can conclude that adding a single additional factor does not hamper learning when no excessive variability or perturbation is introduced. This suggests that existing models of reward-based learning can be extended to a multidimensional task with parallel learning and exploration of the different factors.

As participants only received feedback on a single factor in the first phase, it is not surprising that the factorized feedback hampered the reduction of the combined error in the first phase. More surprising is that there seems to be no learning at all; the combined error even tended to increase (Figure 2). The cause is visible in the figure: an increase in the task-irrelevant error. Errors in the task-irrelevant factor can increase because participants will have variability in the planning of both factors. Without feedback, such errors which will lead to a random walk in performance, leading not only to variability, but also to a systematic error ^19,20^.

Although the results show that two factors can be learned in parallel, the number of factors that can be learned in parallel may be limited. First, exploration adds variability to the motor output. If all factors would be explored by the same amount, the added variability increases with the number of factors that are being explored. This conflicts with the general tendency of the motor system to minimize variability ^21,22^. Second, reward-based motor learning may be an explicit process that depends on working memory ^9,23^. When fewer factors are explored, a longer history of performance can be retained in working memory, providing more information for learning.

As a strategy to reduce exploratory variability in multi-dimensional tasks, participants may determine which factor is most task-relevant and explore this factor before the other factors. Such sequential exploration requires that participants solve an attribution problem, which can be resolved quickly according to the literature. For instance, in a 2D trajectory learning task, participants learned the factor (curvature or direction) that was weighted more heavily in the reward function more rapidly ^13^. We however only found mixed evidence for a relation between task relevance and exploration. Exploration in slant depended on task relevance whereas exploration in length did not. One reason why we found inconsistent results for slant and length may be that different strategies were used in the learning of these factors. For length, some participants reported on the exit interview that they counted the duration of their movement in order to control length more precisely. For slant no such strategies were reported. Also, as perception of depth affects both perception of slant and length, the two factors may have been correlated. Therefore, the influence of task relevance on exploration of length may have been hidden by exploration of slant. Moreover, task-relevant exploration has been reported in the literature. In one experiment for instance, variability in velocity increased during adaptation to a velocity dependent force field whereas variability in position increased following adaptation to a position-dependent force field ^24^.

As a strategy to reduce working memory load in multi-dimensional tasks, participants may rely less on explicit forms of learning when a large number of factors is task-relevant. Our exit interview revealed no difference in explicit learning between groups but explicit learning can be studied more sensitively using for instance a double-task paradigm ^9^, by asking participants to report explicit strategies ^25^ by limiting movement preparation time ^26^ or by instructing participants to drop any strategy they used to score more points in the retention phase ^23^.

## Methods

### Participants

Participants were 60 students of the Vrije Universiteit Amsterdam (age 22.3 ± 3.8; 20 male, 38 female, 2 unregistered sex; 48 right handed, 7 left handed, 5 unregistered handedness). Participants had adequate stereovision (acuity < 60”) as assessed with the StereoFly test and adequate eye sight in our set-up as assessed by asking them to read aloud a text simulated at a distance of 50 cm and in a font size of 4 cm. We used a between-participants design in which participants were, in a random order, assigned to one of three feedback factorization groups. The Slant First and Length First groups received factorized feedback whereas the Combined group received combined feedback. Ethical approval for the study was provided by the local ethical committee (VCWE) of the Vrije Universiteit Amsterdam in accordance with the declaration of Helsinki. Participants provided informed consent prior to participating in the study.

### Set-up

We used an HTC Vive virtual reality set to generate stimuli and record movements. The movement task was performed with a controller that participants held in their dominant hand. We simulated a simple virtual environment (Figure 4a) in which participants stood behind a pole (height 80% of headset height above the floor) with a red ‘starting sphere’ (diameter 6 cm) on top. The visual target was a 16 cm line (1 cm width) with its center 5 centimeters behind the starting sphere, slanted 47° backwards around the fronto-parallel axis. In order to facilitate moving towards the starting sphere, a white 5 cm diameter ball could provide feedback of hand position. Above the target, the trial number and the cumulative score were displayed.

**Figure 4.**
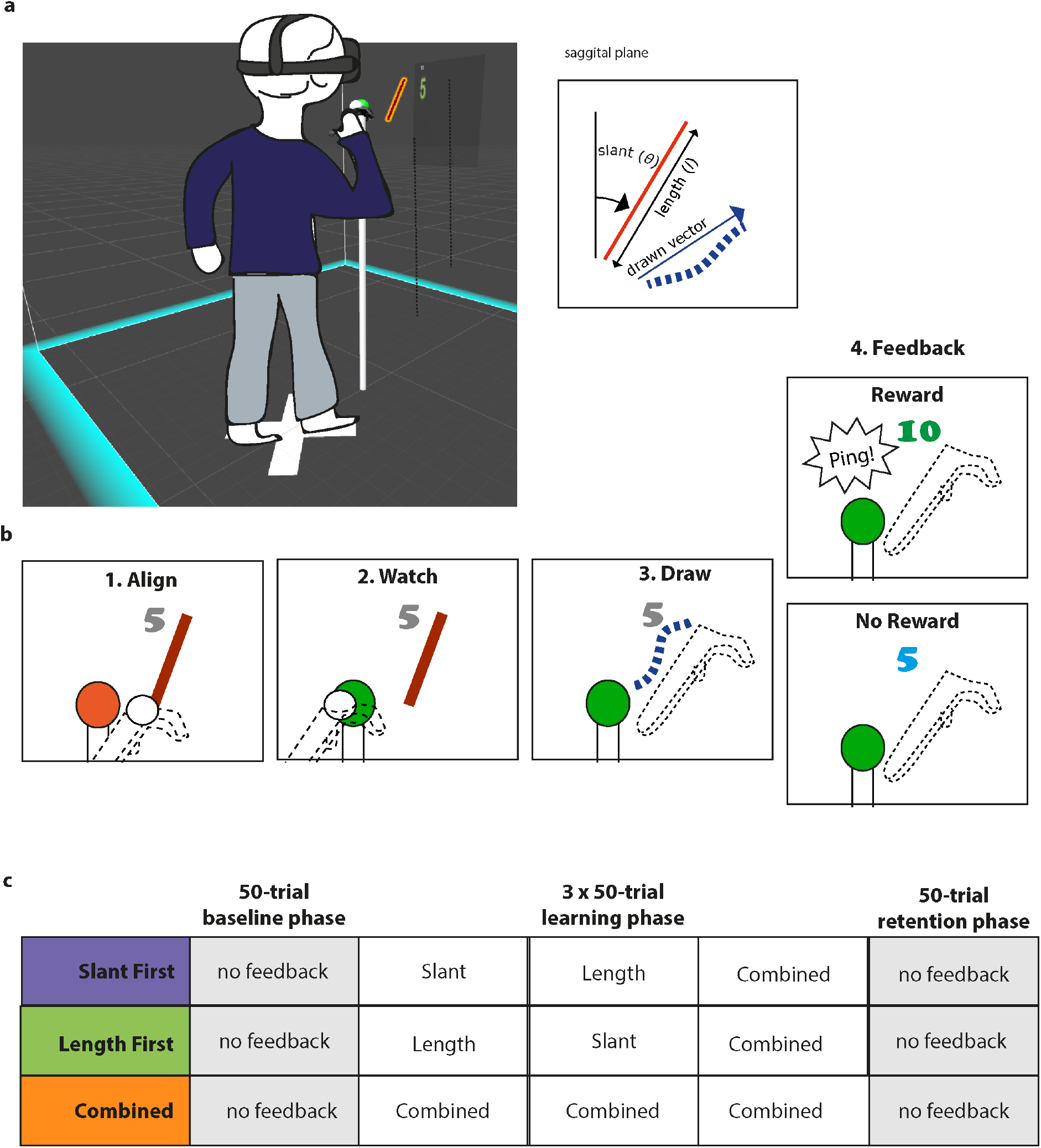
Methods. **a)** Virtual environment with the starting sphere and line target. **b**) Steps within a trial. When a trial was rewarded, the score turned green and 5 points were added to the cumulative score. Otherwise, the score turned blue. **c)** Experimental groups (Slant First, Length First, Combined) and experimental phases.

### Task

Motor learning was assessed in a 3D trajectory matching task in which participants were asked to repetitively copy a remembered line by moving the hand-held controller. To start a trial (Figure 4b), participants were asked to align the controller with the starting sphere. To help them do so, visual feedback on controller position was rendered within a 10-centimeter radius of the starting sphere. When they had successfully done so, the feedback on controller position disappeared, the starting sphere colored green and the target line was visible for 500 ms. After the target had disappeared, the participants were to move the controller to one endpoint of the remembered rod, pull the trigger of the controller, move in a straight line to the other endpoint, and release the trigger when finished – a movement akin to drawing with a pen. Controller vibration provided haptic feedback that a copying movement was recorded. If the controller left the starting sphere too early, the starting sphere colored red and the controller had to be returned before a copying movement could be recorded. In this way, we ensured that participants were copying a remembered line, so that visuo-proprioceptive matching errors ^27^ would not affect task performance. Once the trigger was released, visual feedback on progress and, depending on the experimental phase, performance was provided. After a 300 ms intertrial interval, the participant could initiate the next trial by aligning the controller with the starting sphere.

The experimental phase determined the type of feedback that was provided (Figure 4c) In a 50-trial baseline phase, the trial number was updated, but no reward feedback was provided. In the subsequent three learning phases of 50 trials each, performance feedback was provided based on a ‘drawn vector’ between the starting and endpoint of the drawn line in the sagittal plane. After these three learning phases, participants performed a 50-trial retention phase without any performance feedback. Two factors of the drawn vector could contribute to the performance feedback during the learning phases: slant and length (Figure 4a). Between groups we varied whether these factors were trained sequentially in the first two learning phases (‘factorized’ training) or in a combined manner. The slant factor was ‘vertical’ slant in the sagittal plane, the length factor was the vector length and the feedback on the combined factor was determined based on the vector difference between the drawn vector and the target line.

Performance feedback was provided according to an adaptive success criterion in which trials were rewarded when the amplitude of the relevant error was smaller than a median of the last 5 trials or when the error in the combined factor was smaller than 2 cm:

Participants in the Length First group performed the first learning phase with feedback based on the length, the second learning phase with feedback based on the slant and the third learning phase with feedback based on the combined factor. Participants in the Slant First group performed the first learning phase with feedback based on slant, the second learning phase with feedback based on length and the third learning phase with feedback based on the combined factor. Participants in the Combined group performed all three learning phases with feedback based on the combined factor.

### Procedure

Prior to the experiment, we measured eye distance with a ruler and stereovision with the StereoFly test. Next, participants received visual and auditory instructions on the experimental task. All participants were instructed that they should try to match the target line as accurately as possible with the movement of the controller. The Slant First group was told that their scores would first depend on slant, next on length and finally on the combination of the two. The Length First group was told that their scores would first depend on length, next on slant and finally on the combination of the two. Participants in the Combined group were told that their scores would depend on the combination of slant and length. Illustrations were used to inform the participants how slant and length were defined.

After the instructions, participants put on the headset. We checked visual acuity by asking them to read aloud a participant code simulated at a 50 cm distance and a character size of four cm. We let participants familiarize themselves with the drawing task in four practice trials in which a different target trajectory was shown and, in contrast to all phases of the experiment, full visual feedback on the drawn trajectory was provided. After that, the experiment started.

As motivation may influence how participants learn from score rewards, motivation was assessed after the baseline phase and learning phases of the experiment using a Quick Motivation Index ^28^ in which participants responded orally to the following two questions that were posed by the experimenter using a 1-10 numerical scale: “How much did you enjoy the task until now?” and “How motivated are you to continue?”.

When all five phases of the experiment were finished, the participants’ total score was attached to the scoreboard. After the experiment, participants completed an exit interview in which we asked them about handedness, age, sex, height, clarity of the instructions, explicit knowledge of performance errors and strategies to score points.

### Data analysis

We used three types of error in our analysis: slant error, length error and combined error. For comparison between factors all errors were expressed in centimeters (signed scalars). To express the slant error (*e_θ_*) in centimeters, we used the length of the target line (*l_target_*), the vertical slant of the drawn vector (*θ_drawn_*) and the vertical slant of the target line (*θ_target_*)

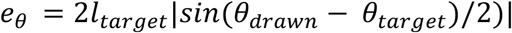

Errors for which *θ* was smaller than the target angle were defined negative, whereas other values were defined positive. To express the error in the combined factor as a scalar representing changes in the direction of interest, we calculated the combined error by projecting the vector error onto the median vector error in the last 20 trials of the baseline phase. In earlier publications we referred to this error as the primary error ^29–31^.

For comparison across participants, the slant error, length errors and the combined error were normalized by the baseline bias such that a value of 1 represented no learning whereas a value of 0 represented complete learning. Negative values would indicate overcompensation. We determined the baseline bias as the median error in the last 20 trials of the baseline phase. We used only the last 20 trials because some participants showed significant drift during the baseline phase. As a measure of learning within a phase, we will report the *gain:* 1 – the median normalized error in the last 20 trials of the phase.

Exploration was analyzed based on a measure of trial-by-trial variability ^14^ in which the amplitude of changes in the slant error (*Δ_θ_*), in the length error (*Δ_l_*) and in the combined error (*Δ_y_*) between trial *N* and trial *N+1* were calculated. Trial-by-trial variation in a phase was the median *Δ* within a phase.

### Statistical tests

Visual inspection of the data (Figure 2 and Figure 3) showed that for some groups the data did not follow a normal distribution. Therefore non-parametric tests were used to test the predictions. Our first prediction was that the learning of an individual factor (slant or length) is fastest when it is the only task-relevant factor. We therefore predicted that for a single factor (slant or length) the gain at the end of a phase would be greater if the factor’s task relevance would be greater. Thus: in the first learning phase, gain of slant would be greater for the Slant First group than for the Combined group and gain of length would be greater for the Length First group than for the Combined group. These predictions were tested using Mann-Whitney U tests.

Our second prediction was that learning of both factors (slant and length) is faster when they are practiced sequentially. We therefore predicted that gain of the combined error in the third learning phase would be greater in the Slant First and Length First groups compared to the Combined group. Again, Mann Whitney U tests were used to test these predictions.

As exploration is the basis of reward-based learning we also assessed how the exploration depended on the factorization. We therefore performed a linear regression of the exploration in a factor on the task relevance and phase number. When the reward did not depend on the factor, we used a value of 0 for the task relevance, when the reward depended only on the factor, we used a value of 1 and when the reward depended on the combined error, we used a value of 0.5.

## Data availability

The datasets generated during and analyzed during the current study and the MatLab code used in the analysis are available in the Open Science Foundation repository, [https://osf.io/vsdt5/].

## Acknowledgments

We thank Duncan Brewster for measuring participants.

## Author contributions

KK designed the experiment, wrote the manuscript and analyzed the data

NM tested participants and wrote the manuscript

JBJ wrote the manuscript

## Additional Information

The authors have no competing interests to declare.

